# Characterization of Collaborative Cross mouse founder strain CAST/EiJ as a novel model for lethal COVID-19

**DOI:** 10.1101/2024.04.26.591329

**Authors:** Candice N. Baker, Debra Duso, Nagarama Kothapalli, Tricia Hart, Sean Casey, Tres Cookenham, Larry Kummer, Janine Hvizdos, Kathleen Lanzer, Purva Vats, Priya Shanbhag, Isaac Bell, Mike Tighe, Kelsey Travis, Frank Szaba, Olivia Bedard, Natalie Oberding, Jerrold M. Ward, Mark D. Adams, Cathleen Lutz, Shelton S. Bradrick, William W. Reiley, Nadia Rosenthal

**Affiliations:** The Jackson Laboratory, Bar Harbor, ME, USA; Trudeau Institute, Saranac Lake, NY, USA; The Jackson Laboratory, Farmington, CT, USA; National Heart and Lung Institute, Imperial College London, UK

## Abstract

Mutations in SARS-CoV-2 variants of concern (VOCs) have expanded the viral host range beyond primates, and a limited range of other mammals, to mice, affording the opportunity to exploit genetically diverse mouse panels to model the broad range of responses to infection in patient populations. Here we surveyed responses to VOC infection in genetically diverse Collaborative Cross (CC) founder strains. Infection of wild-derived CC founder strains produced a broad range of viral burden, disease susceptibility and survival, whereas most other strains were resistant to disease despite measurable lung viral titers. In particular, CAST/EiJ, a wild-derived strain, developed high lung viral burdens, more severe lung pathology than seen in other CC strains, and a dysregulated cytokine profile resulting in morbidity and mortality. These inbred mouse strains may serve as a valuable platform to evaluate therapeutic countermeasures against severe COVID-19 and other coronavirus pandemics in the future.

## Introduction

The COVID-19 pandemic caused by SARS-CoV-2 has had tremendous impacts on humanity that are ongoing today, including significant mortality and morbidity. The pandemic demonstrated in real time how a novel respiratory RNA virus, well adapted to infection of humans, can spread rapidly among the global population despite reactive mitigations. Due in part to encroachment on wildlife habitats, there are recurring viral outbreaks around the world that represent significant threats to human health should sustained person-to-person transmission be established^1,2^. The key to relieving the disruptive and deadly consequences of future pandemics is to devise proactive strategies and solutions to halt virus spread at the earliest possible stages. These strategies should be rooted, in part, to the goal of developing broadly effective vaccines and antiviral therapeutics to combat viruses for which we lack prior knowledge.

COVID-19 has been associated with a wide range of outcomes in human infections, ranging from asymptomatic or subclinical disease to hyperinflammation, acute respiratory distress, and death^3,4^. Moreover, the problem of long COVID, well documented yet poorly understood, occurs in a minority but increasing numbers of infections^5^. There are clear risk factors for severe disease that include advanced age, obesity, and being immunocompromised. In addition, immunological markers such as autoantibodies to type I interferons have been correlated with life-threatening SARS-CoV-2 infections^6^. However, these and other clinical factors do not fully explain susceptibility or resistance to severe COVID-19.

Human genomes have been shaped through co-evolution with pathogenic organisms that have imposed selective pressure on human populations. With the advent of modern medicine in resource-rich nations, selection pressures imposed by human pathogens have moderated, though the global burden of infectious diseases remains substantial^7^. Human genetic variation contributes significantly to the spectrum of infection outcomes. For pathogenic human viruses, well documented common host genetic variants have been strongly associated with resistance/susceptibility to human immunodeficiency virus (HIV), hepatitis C virus (HCV), and other viruses^8,9^. Not surprisingly, common host variants impacting viral disease susceptibility are enriched in genes that regulate immune responses, including human leukocyte antigen genes, interferon genes, and transcription factors^10^.

The development of new vaccines and therapeutics for emerging viruses requires well-characterized, robust, and genetically relevant small animal models. Mouse strains exhibit a wide range of disease susceptibility due to genetic variation but, for many viruses, adequate mouse models are lacking. Disease models for infectious diseases that recapitulate essential pathological features observed in human infections are extremely valuable tools for the development of medical countermeasures against emerging viruses. Proactive development of such models for RNA viruses from families known to harbor epidemic or pandemic species would provide a critical head start for rapid testing of candidate anti-virals and vaccines.

During the early days of the COVID-19 pandemic, it was determined that the spike protein of SARS-CoV-2, like that of SARS-CoV, did not efficiently mediate viral entry via the mouse ACE2 receptor (mACE2)^11,12^ due to apparent low affinity between spike protein and mACE2. Thus, productive infection with the ancestral SARS-CoV-2 Wuhan isolate was not observed in any strains of mice tested^13^. To overcome this barrier, researchers turned to using a transgenic mouse that overexpresses human angiotensin converting enzyme 2 (ACE2) driven by the K18 promotor (K18-hACE2 mice), previously used to study SARS-CoV infection^14^. While these mice provided a platform during the early days of the pandemic to test novel therapeutics and vaccines, they did not ultimately recapitulate the human disease or pathology^13,15,16^. Since the K18-hACE2 mice are on a C57BL/B6J (B6J) background it was unclear if other genetically diverse strains might result in a disease phenotype.

In a previous study^17^ we assessed the impact of host genetics on immune responses and COVID-19 severity by creating a panel of genetically diverse mice that accurately modeled the highly variable human response to SARS-CoV-2 infection. We exploited the Collaborative Cross (CC), a genetically diverse compilation of mice which has been used to identify a number of genetic-based host susceptibility factors to infections^18^. Hybrid offspring generated by crossing eight founders of the CC with inbred mice carrying a transgene expressing human angiotensin converting enzyme 2 (K18-hACE2) were infected with the SARS-CoV-2 Wuhan strain. The resulting spectrum of survival, viral replication kinetics, and immune profiles recapitulated the complexities of SARS-CoV-2 virus replication, dynamics and inflammatory profiles, underscoring the importance of host genetics in interpreting or predicting individual responses to viral infection.

As mutations in SARS-CoV-2 variants of concern (VOCs) have expanded viral host range to mice^19^ we evaluated responses to infection with multiple VOCs in the eight non-transgenic CC founders compared to the canonical K18-hACE2 transgenic mouse model. Notably, all mouse strains were susceptible to infection by VOCs, confirming that mutations accumulated in VOCs rendered them capable of infecting genetically diverse wildtype mouse strains. We observed a range of disease susceptibilities and viral burdens: wild-derived strains (WSB/EiJ, PWK/PhJ and CAST/EiJ), each showed differential disease severity, survival, and viral burden, whereas other CC inbred strains were resistant to disease despite measurable lung viral titers. We further characterized a highly susceptible wild-derived strain, CAST/EiJ, as a novel mouse model for severe COVID-19 that develops high lung viral burdens without spread to the brain, lung pathology, and dysregulated cytokine profile, resulting in morbidity and mortality. We posit that this model will serve as a valuable platform for which to evaluate medical countermeasures against SARS-CoV-2 and other coronaviral pandemics in the future.

## Results

### Genetically diverse mice display varying degrees of direct susceptibility to SARS-CoV-2 VOC infection

The emergence of mutations in the spike protein of SARS-CoV-2 VOC has rendered these viruses capable of engaging the mouse ACE2 receptor and infecting B6J mice^19,20^. To determine the infectivity of other mouse strains, we infected females and males from the CC founders as well as mice heterozygous for the K18-hACE2 transgene (Tg) with 1×10^5^ plaque forming units (PFU) of Beta VOC (hCoV-19/USA/MD-HP01542/2021 Lineage B.1.351) (Figure 1a and b) and examined infected mice for weight loss (Figure 1c and d) and survival (Figure 1e and f). As previously reported, K18-hACE2 mice began to show weight loss starting 2-3 days post infection (dpi) and continued to lose weight until they reached humane endpoints (loss of >25% of their pre-infection body weight or being found moribund) at day 5 to 7 post-infection^17,21^. Commonly used strains, B6J, A/J, NOD scid gamma (NSG), NZO/HlLtJ (NZO), and 129S1/SvlmJ (129S1) mice, were relatively resistant to Beta VOC infection (Figure 1c and d). NSG, created on the NOD/ShiLtJ background from the CC Founders, are immunodeficient and were used in this study to test both the genetic background as well as the immune response to infection. 129S1 and NZO mice exhibited weight loss up to 10-15% and 5-10% of pre-infection weights, respectively, directly after infection. 129S1 mice regained the weight loss from infection after 3 dpi with almost full weight recovery at 14 dpi unlike the NZO mice which were found to have a plateaued weight loss by 5 dpi with no subsequent recovery.

**Figure 1.**
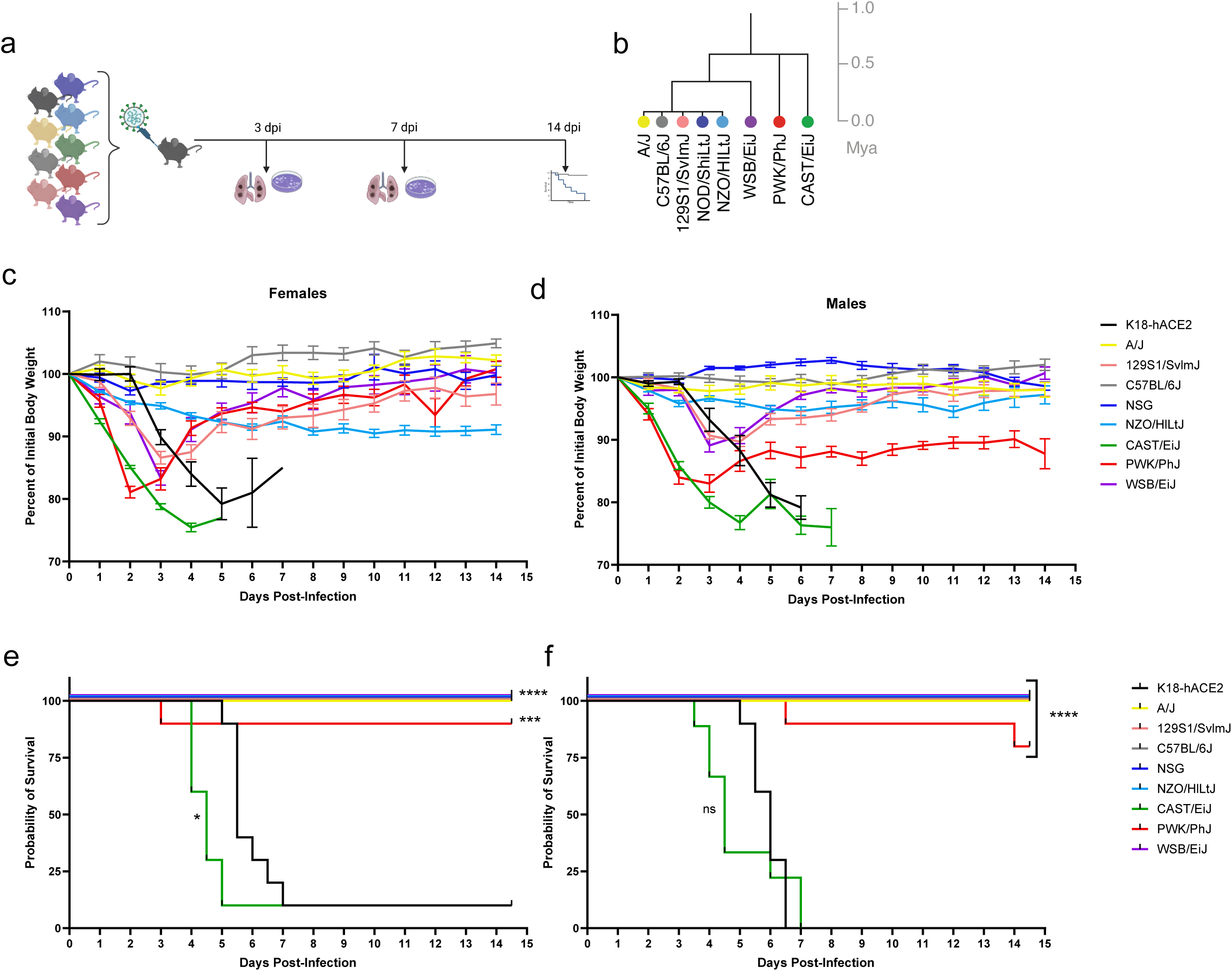
Evaluation of genetically diverse mouse strains for susceptibility to SARS-CoV-2 Beta variant infection in comparison to transgenic K18-hACE2 mice. (a) Experimental schematic showing nine unique mice strains were infected with 1×105 PFU SARS-CoV-2 Beta VOC and assayed at 3 days post-infection (dpi) and 5 dpi for lung viral titer (n=8/strain per time point, 4 males, 4 female) as well as survival (n=20/strain, 10 males, 10 females). (b) Phylogenetic tree showing diverse mouse strains is shown, adapted from Morgan et al. (c-d) Female and male body weights were monitored daily for 14.5 days post-infection. Data are graphed as mean percent of initial body weight ± SEM. (e-f) Probability of survival is shown for each strain. (g) Lung viral titers at 3- and 7 days post-infection graphed as mean titer ± SD.

Compared to the five CC founders representing canonical inbred laboratory strains described above, the three wild-derived inbred CC founders, WSB/EiJ (WSB), PWK/PhJ (PWK), and CAST/EiJ (CAST), were more susceptible to infection with Beta VOC (Figure 1b): all three strains rapidly lost weight (Figure 1c and d). Weight loss was evident at 24 hours of infection in these three strains and continued until day 2.5 post-infection for PWK, day 3.5 post-infection for WSB, and day 5 post-infection for CAST, after which time mice started to regain weight. While a few PWK mice did succumb to infection, occurring sporadically throughout the 14-day study, CAST mice were highly susceptible to lethality and reached humane endpoints after Beta VOC infection, starting on day 4 and ending on day 7 post-infection (Figure 1e and f). To date, CAST is the first non-transgenic strain of mice that has displayed significant lethal susceptibility to SARS-CoV-2 infection.

Viral burdens in the lung tissues were examined in the inbred cohorts of CC founder mice infected with Beta VOC on day 3 and 7 post-infection (Figure 1g). At day 3, K18-hACE2 mice had 5.07±0.37 log_10_PFU of virus in the lungs with similar (not significantly different) levels being detected in 129 mice. B6J mice had slightly lower mean viral burden, 4.72±0.42 log_10_PFU of virus, in the lungs on day 3. In A/J, NSG, and NZO significantly lower viral titers were observed on study day 3 (*p*<0.0001). For WSB mice, high viral burdens in the lungs were comparable to K18-hACE2 mice at 3 dpi (Figure 1g) (mean viral burden of 5.15±0.56 log_10_PFU of virus). With the high susceptibility of the CAST mice, it was not surprising to find significant viral burdens, statistically higher than K18-hACE2 mice, with an average of 5.86±0.08 log_10_PFU of virus per lung on day 3 post-infection. In contrast, PWK mice, which lost significant weight and displayed some susceptibility to the infection, had statistically significant (*p*<0.0001) reduced levels of the virus in the lungs (2.84±0.19 log_10_PFU, day 3 post-infection) compared to K18-hACE2 and CAST mice. By day 7 post-infection, only the NSG mice and a single surviving K18-hACE2 mouse had residual infectious virus in the lungs, likely due to delayed viral clearance mediated by immune system (Figure 1g).

Viral dissemination to the brain leading to encephalitis has been cited as a cause of death in the K18-hACE2 mice^22^, a manifestation not observed in human infections. A productive infection in these humanized transgenic mice is predominantly seen in the olfactory system with neuronal death, perivascular hemorrhage, and leukocyte infiltration in the perivascular space and blood vessels. While we could detect virus in the brains of some K18-hACE2 mice on day 3 post-infection with high levels in the only remaining mouse on day 7, none of the founder lines had detectable virus in their brains on day 3 or 7, more accurately mirroring the human tissue tropism of SARS-CoV-2 (Supplemental Figure 1a).

To characterize SARS-CoV-2 tropism for varied mouse tissues, we determined levels of viral genomic and subgenomic RNA (sgRNA) in the following tissues and samples; fecal material, heart, liver, and spleen at 3 days post-infection (Supplemental Figure 2a and b). The presence of sgRNA is a signal for actively replicating viral RNA while genomic RNA presence may not reflect active viral replication. Viral genomes were readily detectable at generally variable levels, after normalizing to levels of RNase P RNA, between animals in most tissues examined. CAST and B6J showed high levels of SARS-CoV-2 genomes in liver, while 4 of 6 CAST mice showed undetectable or very low levels in heart tissue. In general, sgRNA levels were lower than genomic viral RNA as expected and similar trends were observed between mouse strains as seen for viral genomes with significant variation between individual animals.

We infected the same mouse lines with another SARS-CoV-2 VOC, Gamma (hCoV-19/Japan/TY7-503/2021 Brazil P.1), which yielded similar results as observed with the Beta variant (Supplemental Figure 3A). Together with data shown in Figure 1, the results showed that most commonly used mouse strains are capable of being infected with SARS-CoV-2 but are highly resistant to disease and can rapidly clear the virus.

### Immunological examination of CAST mice infected with SARS-CoV-2 reveals increased number of macrophages, increased IL-6 expression, and histological damage

To gain insight into what lung cytokines might distinguish the susceptibility of CAST mice to SARS-CoV-2 infection from resistant B6J or be shared with the susceptible K18-hACE2 mice, we quantified a panel of analytes known to be important mediators of the immune response. Lung homogenates from uninfected mice or mice infected with Beta VOC at 1×10^5^ PFU for 1, 3, and 5 days were analyzed for 30 cytokines (Figure 2a). The cytokine signatures of CAST and K18-hACE2 mice show sustained increases at 3- and 5-days post-infection for a number of known pro-inflammatory chemokines and cytokines. CAST mice had elevated and sustained levels of IL-6 present in the lung compared to both B6J and K18-hACE2 (Figure 2b). Similar results were observed for Groα, MCP-5, and IL-1β (Figure 2c-e). Some cytokine signatures were similar between K18-hACE2 and CAST, including MCP-1 and MCP-3 (Figure 2f and g). It is possible that elevated levels of these proteins in CAST mice may result from relatively high viral burdens and drive the immunopathology observed with this strain^23,24^.

**Figure 2.**
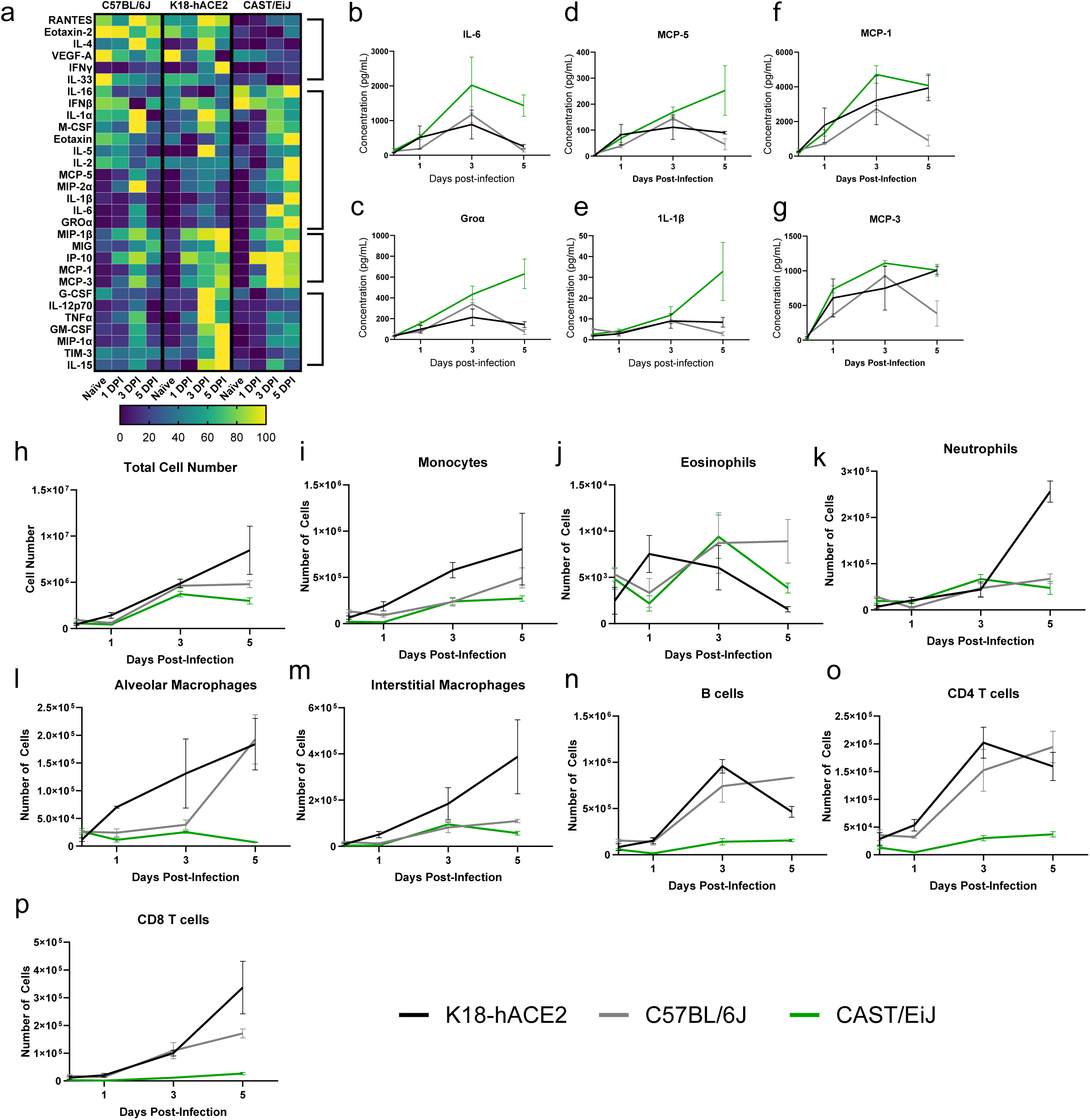
Cytokine and flow cytometric analysis from SARS infected lung tissue show cytokine storm in CAST and K18-hACE2 mice. Mice were infected with 1×105 PFU SARS-CoV-2 Beta VOC. (a) Heatmap of cytokine and chemokine responses from naïve, 1-, 3-, and 5 days post-infection. (b-f) Il-6, Groα, MCP-5, IL-1β, MCP-1, MCP-3 levels in CAST/EiJ (green), C57BL/6J (grey) and K18-hACE2 (black) mice. (h-p) Total number of indicated cell types in lung samples at the indicated timepoint. The data are representative of two experiments of similar design, wherein 5 mice were used per group.

Based on the early timing of the susceptibility of the CAST with high viral burdens it is suggestive that defects in innate immunity could likely be contributing factors towards morbidity. Therefore, we phenotypically analyzed the innate immune cells within the lung tissue of CAST mice after infection with SARS-CoV-2 in comparison to K18-hACE2 and B6J. The frequency and absolute numbers of multiple immune cell types, including B cells, interstitial and alveolar macrophages, neutrophils, CD8 T cells, CD4 T cells, monocytes and eosinophils were analyzed by flow cytometry at baseline (day 0) and days 1, 3, and 5 post-infection (Figure 2h-p and Supplemental Figure 4a-h). The overall number of immune cells in the lung tissues increased over the course of infection for each mouse strain (Figure 2h), consistent with active SARS-CoV-2 infection. Analysis of numbers of specific cell types revealed some similarities and apparent differences between mouse strains. Since K18-hACE2 mice are on a B6J background it was not surprising that the development, recruitment and magnitude of the adaptive immune response, B cells and T cells, was similar between these two strains (Figure 2n, o and p cell number; Supplemental Figure 4f-h, frequency). Yet some differences were seen between these strains starting on day 3 where K18-hACE2 mice had an increased recruitment of monocytes, neutrophils, and interstitial macrophages (Figure 2l, k, and m) which likely is a result of the increased lung damage in the susceptible K18-hACE2 strain compared to the resistant B6J strain. CAST lungs did show an increase in the total cell number albeit lower cellularity compared to B6J or K18-hACE2 mice at all time points examined but did not show increases in the numbers of alveolar macrophages over the course of infection in contrast to B6J or K18-hACE2 (Figure 2l). In general, the total cellularity in CAST mice did not show an obvious change within a specific cell population. The most notable change in a cellular population found in the CAST mice was an increase in the frequency but not the total cell number of interstitial macrophages, increasing over the course of infection, which was not seen in B6J or K18-hACE2 mice (Supplemental Figure 4e). These data show that the cellular profile of the immune cell response between B6J and K18-hACE2 mice were similar compared to the CAST mice. Yet the differences seen in cellularity were not as apparent when frequencies were examined between these 3 strains. This in part could be due to the overall animal size difference and tissue weight differences that exist between the B6J and K18-hACE2 strains versus the CAST strain of mice.

### Susceptibility of CAST mice to varying doses of SARS-CoV-2

To further investigate the lethality seen in CAST mice, we next examined if survival was dependent on the viral dose administered. Mice were infected with 1×10^5^ PFU, 1×10^4^ PFU, or 1×10^3^ PFU of SARS-CoV-2 Beta VOC. Female and male mice were weighed for 14 dpi (Supplemental Figure 5a and b) and monitored for survival after infection (Supplemental Figure 5c and d). Females at all dosages lost weight precipitously and none survived at any dose (Supplemental Figure 5a and 5c). Males at the lower dosages (1×10^4^ PFU, and 1×10^3^ PFU) lost weight as early as day 1 post-infection and continued until day 4 post infection where surviving mice started to regain weight slowly but did not fully recover their initial starting weight even after 14 days post-infection (Supplemental Figure 5b and 5d). In contrast to clinical and preclinical findings of sex bias in SARS-CoV-2 susceptibility, CAST male mice had an increased survival rate compared to female mice^25^. This effect was only seen when the viral inoculum was lowered to 1×10^4^ PFU or 1×10^3^ PFU per mouse (Supplemental Figure 5c and d) suggesting that lower viral titers might help reveal more subtle but important aspects of infection in this strain.

### Histological analysis of lung tissue from infected mice

To understand the cellular complexity and composition of the CAST, B6J and K18-hACE2 mice, we examined if lung histological differences were observed between these strains using SARS-CoV-2 viral nuclear protein (NP) (Figure 3a), hematoxylin and eosin (Figure 3b) as well as an antibody specific for ACE2 expression (Figure 3c). All three strains showed expression of SARS-CoV-2 viral nuclear protein (NP) antigen in the bronchiolar club cells (Figure 3a, blue arrows and d). Unlike B6J mice, K18-hACE2 mice also displayed high viral antigen on day 1 post-infection in alveolar type 1 pneumocytes (AT1 cells) with a few alveolar type 2 pneumocytes (AT2 cells) staining positive (Figure 3a, green and blue arrows, respectively). Uniquely, CAST mice only showed positive NP antigen in AT2 cells, not in AT1 cells. While differences existed in the localization of viral antigen between all 3 strains, both B6J and CAST mice had the highest NP IHC staining in club cells on day 1. This was eclipsed on day 3 and 5 where K18 hACE2 mice displayed the highest level of staining. Not surprisingly, based on the previous plaque data, B6J mice had little detectable viral antigen staining by IHC on day 3 and day 5. CAST mice retained high levels of viral antigen staining residing in AT2 cells and to a lesser extent in the bronchiolar club cells at later time points.

**Figure 3.**
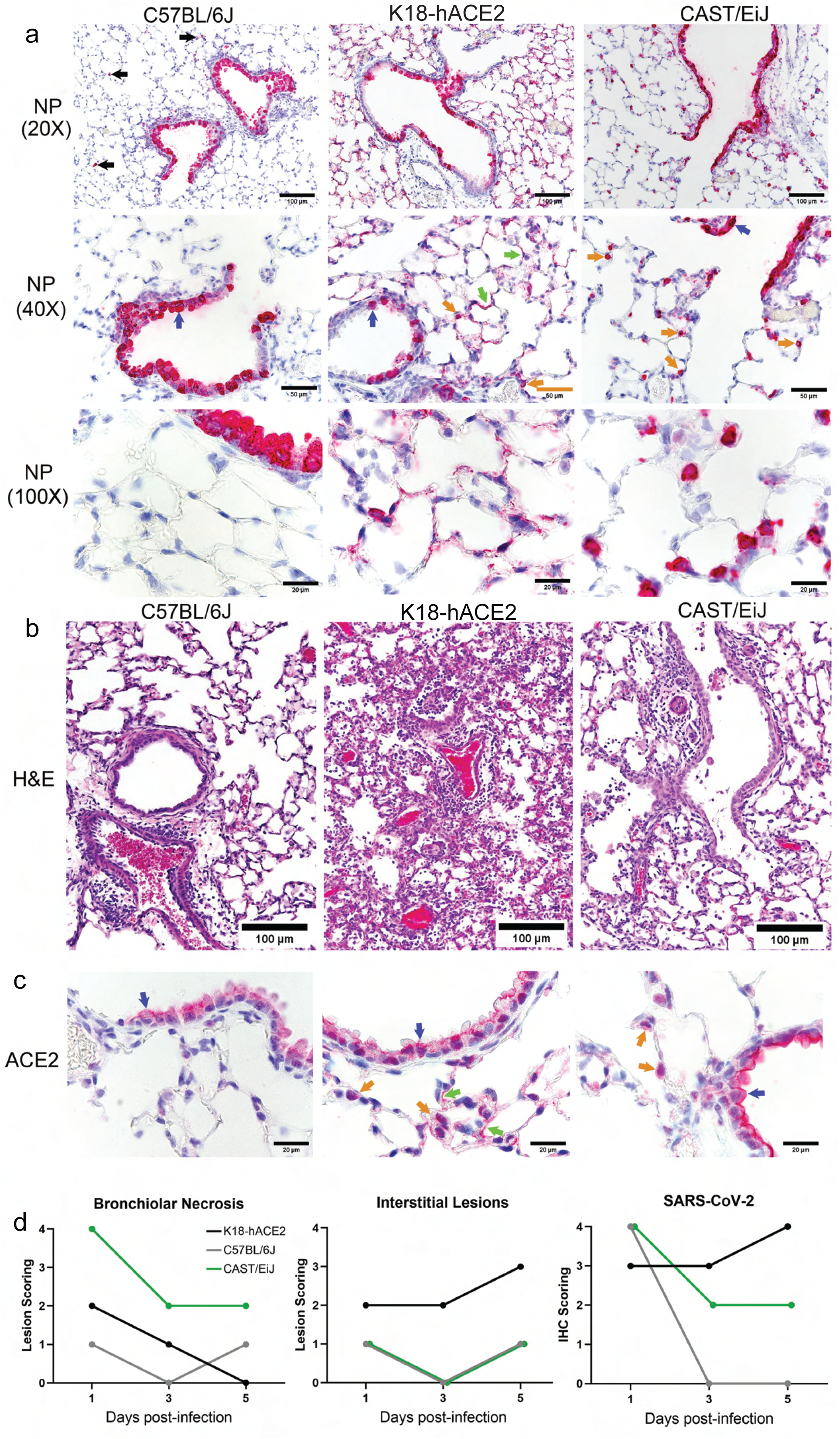
Histological and immunohistoche mical analysis of SARS-CoV-2 infected lung tissues. (a) a-NP antibody staining of lung sections was performed to identify infected cells (in red) in the lungs of each mouse strain at day 3 post-infection. Black arrows point to low frequency infected macrophages found in C57BL/6J mice. Blue arrows point to infected bronchiolar club cells. Green arrows point to infected AT1 cells and orange arrows point to infected probable AT2 cells. (b) Hemotoxin and eosin-stained lung sections from each

Perivascular cuffing was seen in all strains on day 5 post infection which agrees with increased cellularity seen in the flow cytometry results (Figure 3b). However, both K18-hACE2 and CAST mice had prominent alveolar lesions not seen in B6J mice. CAST mice showed endothelialitis in small vessels and a mild multi-focal interstitial increase of macrophages and/or AT2 cells. CAST mice displayed the highest levels of bronchiolar necrosis throughout all 5 days of infection analyzed (Figure 3d). Yet, possibly due to the expression difference of ACE2 by the K18 promoter, the K18-hACE2 strain of mice displayed the highest levels (most severe) of the interstitial lesions (Figure 3b and d). No brain lesions or viral antigen were seen in mice strains except for K18-hACE2 mice (data not shown) which is not unexpected based on viral plaque results.

The ACE2 receptor antigen was found by immunohistochemistry in bronchiolar club cells of naïve mice of all strains tested and in many AT1 cells and lesser numbers of AT2 cells of K18-hACE2 mice and in many AT2 cells of CAST mice (Figure 3c). The expression pattern of ACE2 correlated with the expression patterns of viral antigen seen after infection.

### Therapeutic intervention of SARS-CoV-2 infection of CAST mice

Based on the accumulated data we reasoned that CAST mice could serve as a useful model for analysis of anti-viral therapeutics. To address this, we infected K18-hACE2 and CAST mice with 2×10^4^ PFU of Beta VOC and 1 day post infection, mice were treated with a protease inhibitor (PF-1332) at 500 mg/kg/day or vehicle for the first 4 days of infection. K18-hACE2 mice that survived with PF-1332 treatment nearly regained their starting body weight, while treated CAST mice lost weight and generally did not gain weight by 14 days post-infection (Figure 4a). Treatment with PF-1332 significantly enhanced survival for both mouse strains, leading to 40% and 70% survival for CAST and K18-hACE2 mice, respectively (Figure 4b). Together, these data highlight the usefulness of using CAST mice to evaluate antiviral therapeutic countermeasures.

**Figure 4.**
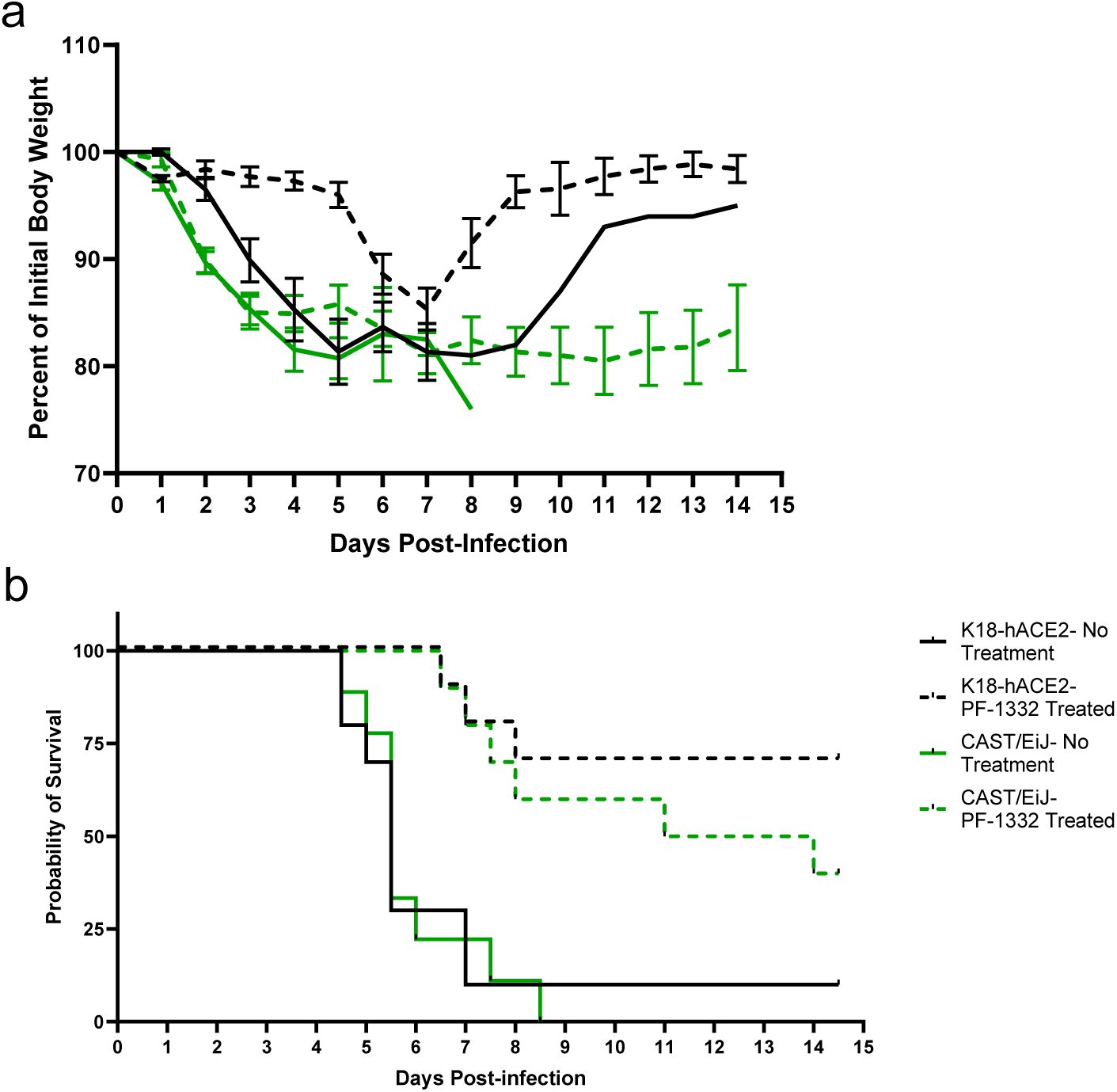
Therapeutic treatment of K18-hACE2 and CAST/EiJ mice infected with SARS-CoV-2 Beta VOC. (a) Mean weights ± SEM for each group over the course of 14.5 days. (b) Survival curves for the indicated groups with or without treatment with PF-07321332 at 500 mg/kg/day. The data are representative of two experiments of similar design, where in 5 mice were used per group.

## Discussion

Human genetic variation has major impacts on infectious disease risk and severity. Prominent examples of this are the unequivocal role of *IFNL3* gene polymorphisms in determining frequency of spontaneous and therapeutic HCV clearance^26,27^ and the strongly protective effect of the *CCR5*Δ32 deletion in HIV infection^28^. For SARS-CoV-2, the human genetic component of disease risk is complex and complicated by the diverse manifestations of infection, ranging from asymptomatic to lethal^29^. While the role of genetics in human disease caused by RNA viruses is well established, the use of genetically diverse mice for establishing appropriate *in vivo* models of disease and understanding relevant host genes that influence susceptibility has been examined for only a limited number of viruses. Notably, specific host genes in mice have been primarily investigated using reverse genetic approaches by evaluating the effects of ablation or humanization of specific genes, such as innate immunity genes^30^. While these studies have yielded valuable information, they do not take into account the simultaneous contribution of genetic variants impacting multiple pathways that may influence infection outcomes.

The inherent ability of a given virus to cause disease in a mouse model that faithfully recapitulates the disease in humans is influenced by multiple factors. First, an RNA virus must have the capacity to replicate and spread in mouse cells. Essential host dependency factors that allow for virus entry, translation, RNA replication, and assembly must be present in the proper levels and cell types for successful virus infection and dissemination. Relatedly, the virus must be able to appropriately cope with host restriction factors and immune responses to cause disease. All these factors may be influenced by host genetic variation. Thus, future studies using mouse models that reflect the genetic diversity of humans are needed both to gain biological insights and to produce novel mouse models of disease which can be used as tools for development of medical countermeasures.

Here, we analyzed the ability of SARS-CoV-2 Beta and Gamma VOCs to cause disease in genetically distinct mice representing the genetic backgrounds of the Collaborative Cross (CC) founder strains compared to a transgenic strain expressing human ACE2 driven by K18 promoter (on a B6J background). Most strains infected with either Beta or Gamma VOC, were highly resistant to infection, with little to no weight loss and uniform survival after challenge despite measurable lung virus titers at 3 days post-infection. This indicates that productive virus infection of the lung is not sufficient to induce a disease phenotype in many of the strains of mice examined. Strikingly, infection of the “wild-derived” strains among CC founders, WSB, PWK and CAST, induced variable disease phenotypes for each strain. Infection of the CAST and PWK mice led to rapid weight loss which was more profound and longer lasting for CAST. Ultimately, all female and nearly all male CAST mice succumbed to infection with Beta VOC by either 5- or 7-days post infection for females and males, respectively. Despite rapid weight loss, PWK mice quickly recovered and showed overall high survival. In contrast, WSB mice showed relatively mild and transient weight loss, initially mirroring that seen with K18-hACE2 mice before exhibiting a rebound at day 4 post-infection. Interestingly, these phenotypes do not correlate with day 3 lung virus titers, as infected PWK mice had >100-fold lower virus titers than other susceptible strains, while CAST mice uniformly had the highest lung titers. Importantly, none of the “wild-derived” CC mouse strains founders had detectable infectious virus in the brain in contrast to K18-hACE2 mice, indicating that disease and lethality in the CAST mice was not due to encephalitis. The susceptibility to mortality observed in the CAST and PWK strains could be explained by their common ancestral background which diverged approximately 0.5 million years ago (Mya) in these two strains (Figure 1B). WSB mice were segregated more recently and were also susceptible to weight loss (but not mortality), whereas B6J, A/J, NOD/ShiLtJ (NOD), NZO, and 129S1 are all quite similar with respect to their phylogenetic background. Thus, the disease phenotypes in these CC strains may more authentically mimic that seen in humans where brain infection has not been reported.

CC strains have been a useful tool to relate genotype to phenotype in mice for many disease states, infectious and otherwise. For pathogenic human RNA viruses, CC strains have enabled understanding of host genetics impacting a variety of viruses, including SARS-CoV^31,32,33,34,35,36^, influenza virus^37,38,39,40,41,42^, Ebola virus^43,44^, West Nile virus^45,46,47,48^, Theiler’s murine encephalitis virus^49,50,51,52,53^, lymphocytic choriomeningitis virus^54^, Zika virus^55^, SARS-CoV-2^17,36^, Rift Valley fever virus^56^, Powassan virus^57^, and recently, Norway rat hepacivirus^58^, which is closely related to HCV. These studies have revealed mouse genetic background impacts overall susceptibility to infection, viral burdens, host survival, levels of tissue inflammation, innate and adaptive immune responses, and differential gene expression after virus infection. More importantly, infection of genetically diverse mouse panels has allowed identification of specific mouse genetic loci responsible for observed phenotypes, which can be broadly applicable to multiple related viruses or highly specific to a particular species^57^. In some cases, genes associated with virus susceptibility can be related to findings derived from human genome-wide association studies^36,59^, validating the CC platform as a tool to understand how human genetic variation impacts outcome of virus infections.

The outcome of patients infected with SARS-CoV-2 has been directly linked with the presence and severity of a cytokine storm. The sustained release of pro-inflammatory cytokines and chemokines such as IL-6, Groα, MCP-1, IP-10, TNF-α, IFNγ and IL-1β, among others, play a significant role in disease severity and survival outcomes^60,61^. Here we show prolonged increases in a unique cytokine signature observed in CAST mice. These hyperinflammatory signaling and reduced clearance of cytokines mimic what is observed in clinical patients presenting with severe COVID infection. Additionally, a shift in the compartmentalization from alveolar to interstitial macrophages has been identified in both autopsy of patients succumbing to SARS-CoV-2 virus as well as CAST animals^62,63^. Additionally, the localization of ACE2 receptor predominately within AT2 cells as well as viral RNA nuclear protein (NP) staining correlates with cases of severe COVID-19^60,64,65^.

The use of antiviral medications within the correct timing window has also proved beneficial to survival outcomes of patients. While K18-hACE2 mice have served to examine vaccines and therapeutics, they are not the ideal model owing to the following: brain infection from which the mice succumb, the pattern of infection within the lung, and lack of cytokine dysregulation, none of which mirror clinical manifestations of disease seen in human patients. Since CAST mice display several similarities to human infection, they could potentially serve as a better model for evaluation of next generation therapeutics and vaccines against SARS-CoV-2. Clearly, both K18-hACE2 and CAST mice responded to therapeutic treatment with the protease inhibitor PF-1332. However, where the K18-hACE2 mice regained weight loss due to acute infection, CAST mice did not regain weight. This treatment regime offers the potential to utilize CAST animals to examine if long-COVID symptoms might be explored in the mouse strain.

Our findings reveal a diversity of responses to SARS-CoV-2 infection in genetically distinct, non-humanized mouse strains. Each of the “wild-derived” strains, WSB, PWK and CAST, were productively infected with associated disease phenotypes that are worthy of further study. In particular, our findings highlight the utility of CAST mice as a model for severe COVID-19 that leads to lethality without associated encephalitis. The similarities to human infection bolster the case for the use of CAST animals as a model for severe SARS-CoV-2 and for evaluation of human coronaviruses that will emerge in the future.

## Methods and Materials

### Mouse Strains

All mouse strains (B6.Cg-Tg(K18-ACE2)2Prlmn/J; JR034869, C57BL/6J; JR000664, A/J; JR000646, 129S1/SvlmJ; JR002448, NOD.Cg-*Prkdc^scid^ Il2rg^tm1Wjl^*/SzJ; JR005557, NZO/HlLtJ; JR002105, CAST/EiJ; JR000928, PWK/PhJ; JR003715, WSB/EiJ; JR001145) were provided by The Jackson Laboratory (Bar Harbor, ME) and maintained under specific pathogen–free conditions. All strains used for the study were housed in a biosafety level 3 (BSL3) environment for the duration of these studies. The research protocol was approved by the Institutional Animal Care and Use Committee of the Trudeau Institute, protocol 21–005. Mice were housed in the animal facility of the Trudeau Institute and cared for in accordance with local, state, federal, and institutional policies in a National Institutes of Health American Association for Accreditation of Laboratory Animal Care-accredited facility. Animals were maintained in IVC cages on negatively pressurized Allentown PNC racks that were HEPA filtered and directly vented though the building’s exhaust system. The racks and animal rooms were negatively pressurized. Access to all facilities is controlled electronically by the buildings management systems and restricted to approved users only. The environment, temperature, and humidity within the animal facility room were constantly monitored by the building management system. Temperatures were also monitored and recorded daily in individual animal rooms by animal care staff using electronic thermometers. All temperature set points were within The Guide recommended range of 68–79 °F for mice. Acceptable Institutional daily fluctuations are between 67 and 74 °F with a humidity range of 30–70%. Light cycles in all animal-holding and procedure spaces are controlled on a 12/12 light/dark cycle.

### Viral Strains and Infection

All viruses were obtained from BEI Resources and propagated on Vero E6 cells (VERO C1008, ATCC). Virus stocks were concentrated using Amicon filters and tittered by standard plaque assay using Vero E6/TMPRSS2 cells (provided by Dr. Sean Whelan – Washington University). On study day 0, all mice were infected with 1x 10^5^ PFU of either hCoV-19/USA/MD-HP01542/2021 (Lineage B.1.351; Beta Variant) or hCoV-19/Japan/TY7-503/2021 (Lineage P.1; Gamma Variant) diluted in PBS (Corning) via intranasal instillation in 0.03 mL. All mice were monitored for clinical symptoms and body weight twice daily, every 12 h, from study day 0 to study day 14. Mice were euthanized if they displayed any signs of pain or distress as indicated by the failure to move after stimulation or presentation of inappetence, or if a mouse had > 25% weight loss compared to their pre-infection body weight. Mouse survival comparisons were carried out using GraphPad using Kaplan-Meier and Mantel-Cox test to compare K18-hACE2 versus each of the eight strains.

### Cell culture and Plaque Assay

Vero E6/TMPRSS2 cells were cultured in Eagle’s Minimum Essential Medium (EMEM, Corning) supplemented with 10% FBS (HyClone) at 37°C with 6% CO_2_. The plaque assay was performed on VeroE6-TMPRSS2 (Vero-TMP) cells. Briefly, 2 × 10^5^ cells per well were grown in 24-well plates and infected with serial dilutions of the viruses for 1 hour at 37°C, and subsequently, 0.75 ml of an overlay medium was added. The overlay medium contains a final concentration of 0.6% microcrystalline cellulose in 1x MEM (Gibco). At 3 days post-infection, the cells were fixed with 10% neutral buffered formalin (Sigma) for 1 hour at room temperature before the overlay medium was removed, and the cells were stained with 0.1% crystal violet solution (95% water and 5% ethanol) for 5 minutes. The crystal violet solution was removed, and plates were allowed to air-dry before counting the plaques. Statistical analysis was carried out using GraphPad using one-way ANOVA and Dunnett’s multiple comparisons test to compare K18-hACE2 to each of the eight other strains.

### Cell Isolation and Flow Cytometry

Lung tissue was prepared by coarsely chopping the tissue followed by incubation in a 0.5 mg/mL solution of collagenase D (Roche) and 500 Units of DNase (Sigma Aldrich) for 30 min at 37 °C. Single-cell suspensions were prepared by dispersing the digested tissues through a 70 μm cell strainer. Red blood cells were lysed with ammonium buffered chloride. Lymphocytes were enriched from digested lung tissue by differential centrifugation using a gradient of 40/80% Percoll (GE Healthcare). Finally live cell numbers were determined by counting with trypan blue exclusion.

Single-cell suspensions were incubated with Fc-block (anti-CD16/32) for 15 min on ice followed by staining with antibodies to: CD103 (2E7), CD24 (M1/69), CD64 (X54-5/7.1) (Biolegend); CD335 (29A1.4), CD4 (RM4-5), CD11b (M1/70), Zombie UV, (ThermoFisher); CD19 (1D3), CD8a (53-6.7), CD3 (500A2), I-A/I-E MHC Class II (2G9), Siglec F (E50-2440), Ly6G (1A8), CD11c (N418), Ly6C (AL-21) (BD); for 45 min on ice. Following staining, samples were washed two times with phosphate buffed saline and resuspended in 2% formalin overnight. The following day the fixed samples were washed with phosphate buffered saline and removed from the BSL-3 lab for acquisition. Fixed and stained samples were run on a LSRII flow cytometer (BD Biosciences) and data were analyzed with FlowJo software (BD Biosciences).

### Histology

The mice were euthanized by CO_2_ and selected tissues were fixed with 10% neutral buffered formalin (Sigma, St. Louis, MO, USA) for 24 h. Once fixed, the brain tissue was separated from the skull. The brain and lung tissue were processed into paraffin (Sakura Tissue-Tek VIP processor) blocks. Tissue sections were cut at 5 microns thickness and placed on plus charged slides for better adherence.

For H&E staining, the slides were dewaxed through several changes of Xylenes, rehydrated through decreasing concentrations of ethanol and finally into Distilled water. Slides were then stained using standard hematoxylin and eosin staining protocols. After staining, slides were dehydrated through increasing concentrations of ethanol followed by several changes of xylene. Coverslips were applied using a permanent mounting medium and allowed to dry overnight.

For IHC, antigen retrieval was performed using 1X citrate buffer (BioGenex 10X Citra solution) under high temperature in a steamer and then cooled to room temperature. BloxAll solution (Vector Labs) was used to block any endogenous enzyme activity in the tissues and was followed by 2.5% normal horse serum (Vector Laboratories, Burlingame, CA, USA) for 1 hour. The primary antibody rabbit anti-SARS-CoV-2 N protein (NB100-56576, Novus Biologics) was diluted 1:500 in 2.5% normal horse serum and incubated on tissues overnight. The slides were rinsed with PBS and then incubated with anti-IgG polymer detection solution (Vector Labs) for 30 minutes at room temperature, rinsed in PBS again and developed using alkaline phosphatase substrate (Vector Red, Vector Labs). The slides were rinsed with distilled water and counterstained with hematoxylin.

### Luminex Assay

The superior lung lobe, upper right lobe, of infected mice was harvested and homogenized in Tissue Extraction Reagent I (Invitrogen), containing a Protease Inhibitor Cocktail (Sigma Aldrich) following the manufacturer’s instructions. Samples were spun, filtered and frozen at -80°C. Lung homogenates from animals harvested on day 1, day 3 and day 5 post infection were assessed for cytokines by a Luminex Procarta Plex (Mouse Procarta Plex Mix and Match 34-plex, Thermo Fisher, Austria) following the protocol provided by the manufacturer.

#### Nucleic acid extraction

Total nucleic acids were extracted from tissue homogenates using the MagMax-96 Viral RNA Isolation Kit (Thermo Fisher, Catalog No. AM1836) automated on the KingFisher™ Flex magnetic particle processor (Thermo Fisher, Catalog No. 5400630). In brief, RNA was extracted from 200µl of homogenized tissues following the manufacturer’s instructions and eluted in 200µl of elution buffer.

#### qRT-PCR to estimate SARS-CoV-2 viral concentrations

Quantitative real-time reverse-transcriptase PCR (qRT-PCR) was performed to determine SARS-CoV-2 viral concentrations. Assays targeting subgenomic RNAs for the E and N genes of the viral RNA (sgE and sgN, respectively) were multiplexed in a single reaction well. TaqPath™ COVID-19 Combo Kit (Thermo Fisher, Catalog No. A47814) was used to assess the total SARS-CoV-2 viral RNA in a separate reaction well of 384 plate. The TaqPath™ targets include the orf1ab, N and S genes. An assay targeting the mouse RNaseP gene (Rpp30) was used as an extraction control to confirm successful extraction RNA of mouse tissues.

For the sgRNA reaction well, the 10ul reaction mix consisted of 5ul of RNA, 2.5ul of TaqPath™ 1-Step Master Mix No Rox (Thermo Fisher, Catalog No. A28523), 0.2ul ThermaStop™-RT Additive (Sigma-Aldrich, Catalog No. TSTOPRT-250), 0.5ul of sgE assay mix, 0.5ul of sgN assay mix and 1.3ul nuclease-free water.

For the TaqPath Combo Kit with RNAseP reaction well, the 10ul reaction mix consisted of 5ul of RNA, 2.5ul of TaqPath 1-Step Master Mix No Rox (Thermo Fisher, Catalog No. A28523), 0.5ul of TaqPath COVID-10 Combo Kit assay mix, 0.5ul of Rpp30 assay mix and 1.5ul nuclease-free water.

The details of all the primer and probe sequences are listed in Table 1. The assay mixes were prepared by mixing the forward and reverse primers and probes at the concentrations listed in Table 1 to yield a 20x working stock, with the final assay concentration at 1x. The qRT-PCR assays were performed in a QuantStudio 7™ Flex Real-Time PCR Systems (Applied Biosystems). Thermal cycling was performed at 25°C for 2 minutes, 53°C for 10 minutes for reverse transcription, followed by 95°C for 2 minutes and then 40 cycles of 95°C for 5 seconds and 60°C for 45 seconds.

**Table 1:**
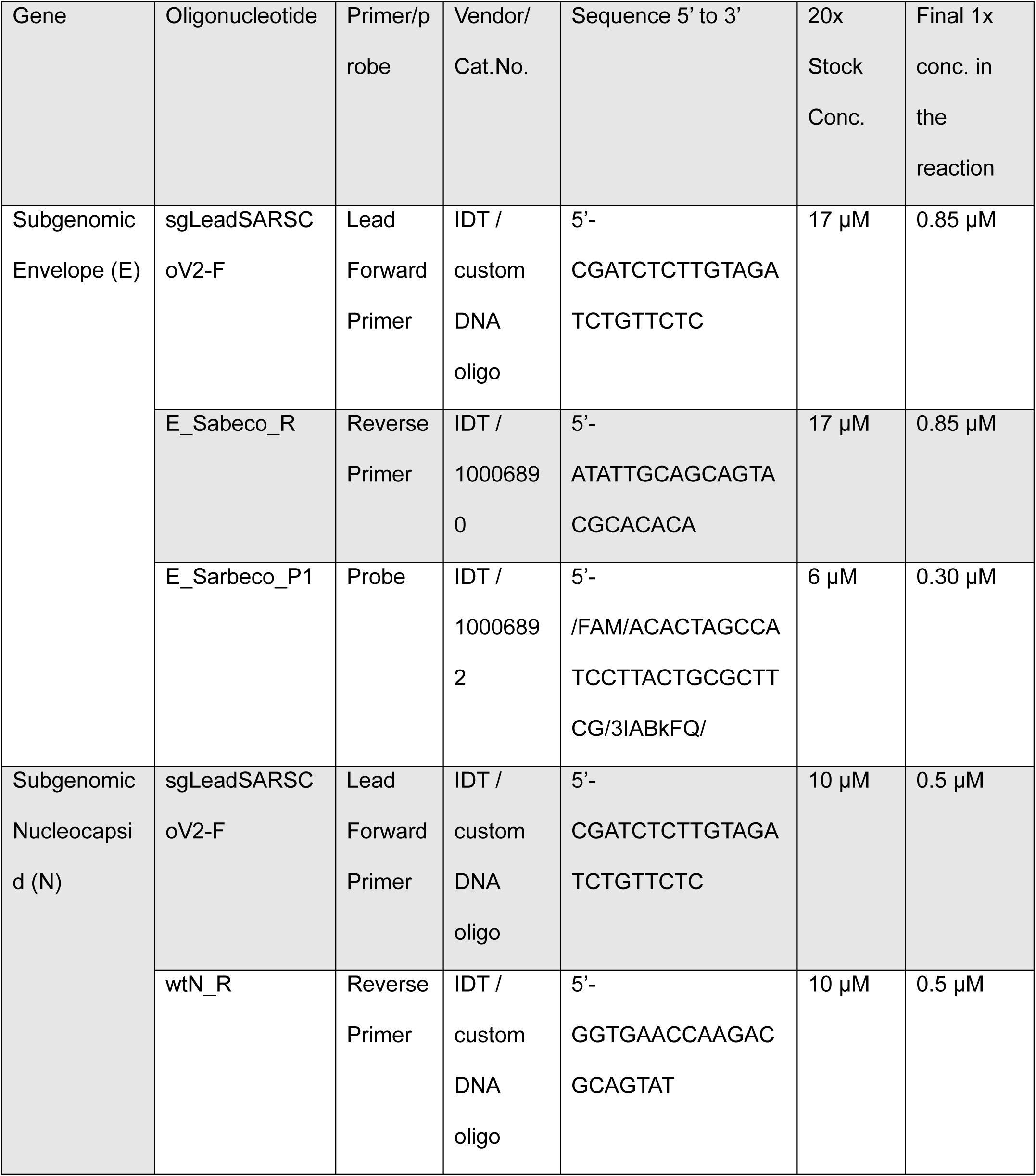
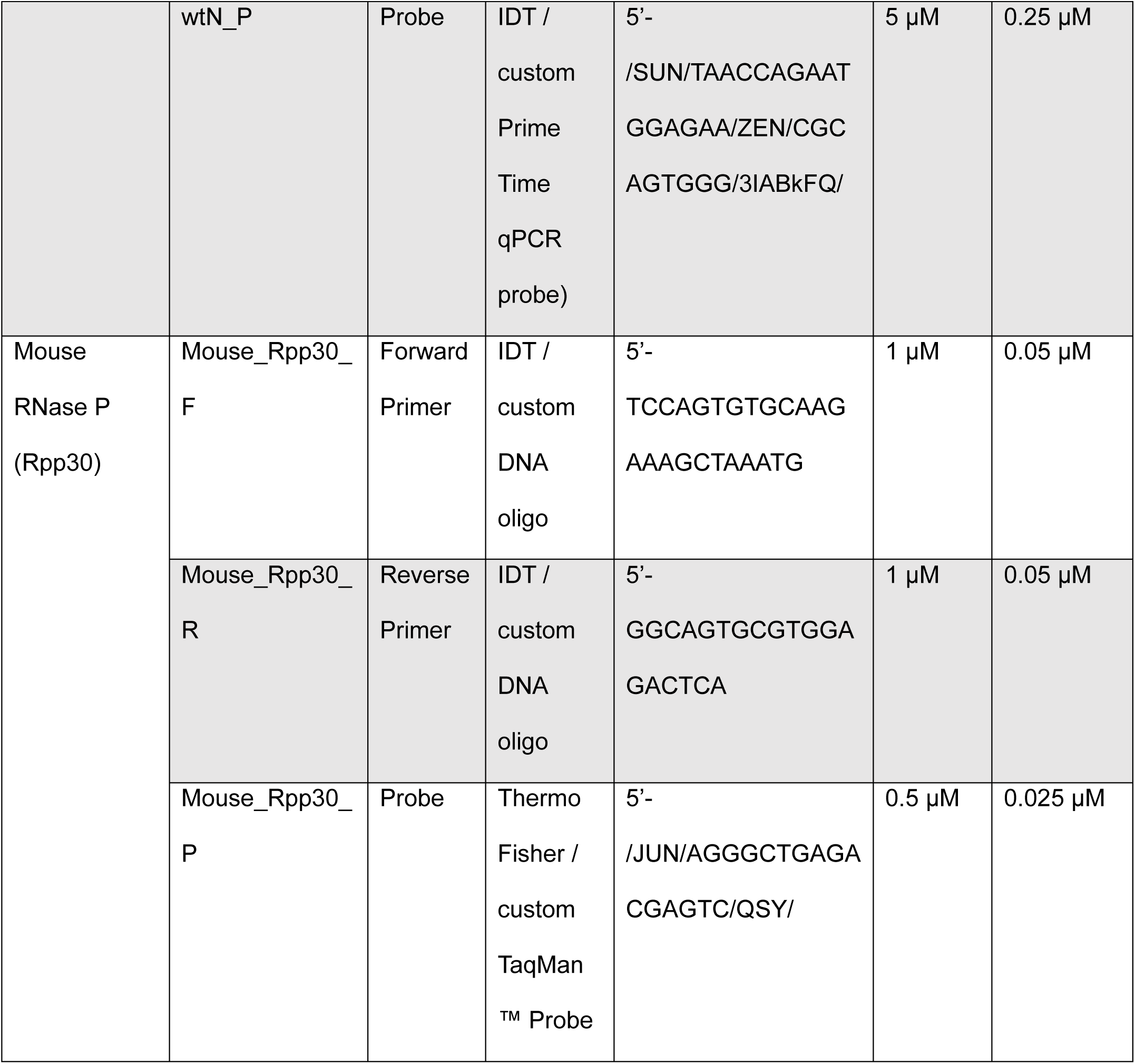
Primers and Probes for RT-PCR.

## Data Availability

The data that support the findings of this study are available from the corresponding authors upon request.

## Acknowledgements

Funding was provided by a grant to the Special Mouse Strain Resources at The Jackson Laboratory from the National Institutes of Health (NIH), Office of Research Infrastructure Programs (P40 OD011102) and by support from the F. M. Kirby Foundation and Anonymous Family Fund to the Trudeau Institute. The following reagent was obtained through BEI Resources, National Institute of Allergy and Infectious Diseases (NIAID), NIH: SARS-Related Coronavirus 2, isolate hCoV-19/USA/MD-HP01542/2021 (Lineage B.1.351), in *Homo sapiens* Lung Adenocarcinoma (Calu-3) Cells, NR-55282, contributed by Andrew S. Pekosz, and SARS-Related Coronavirus 2, Isolate hCoV-19/Japan/TY7-503/2021 (Brazil P.1), NR54982”, contributed by (NIAID). We thank Jeffrey Harder, Daniel Skelly, Steven Munger, and Christopher Norbury for their expert assistance in this publication. Thank you to the animal caretaker staff at both Trudeau Institute and The Jackson Laboratory campuses for their work on these studies.

## Author contributions

All authors reviewed the final manuscript. Viral infections and mouse work were performed by D.D., N.K., T.H., S.C., T.C., L.K., J.H., K.L., P.V., P.S., I.B., M.T., K.T., F.S., O.B., N.D, J.W., M.D.A., and W.W.R. Visualizations were created by C.N.B, S.S.B., J.M.W., W.W.R. Conceptualization of project by: N.R., C.L., and W.W.R. Writing of manuscript by C.N.B, S.S.B, W.W.R, N.K., and N.R. Funding provided by C.L., N.R., and W.W.R.

## Ethics declarations

The authors declare no competing interests.

