## Supplemental Figures for "Characterization of Collaborative Cross mouse founder strain CAST/EiJ as a novel model for lethal COVID-19"

a

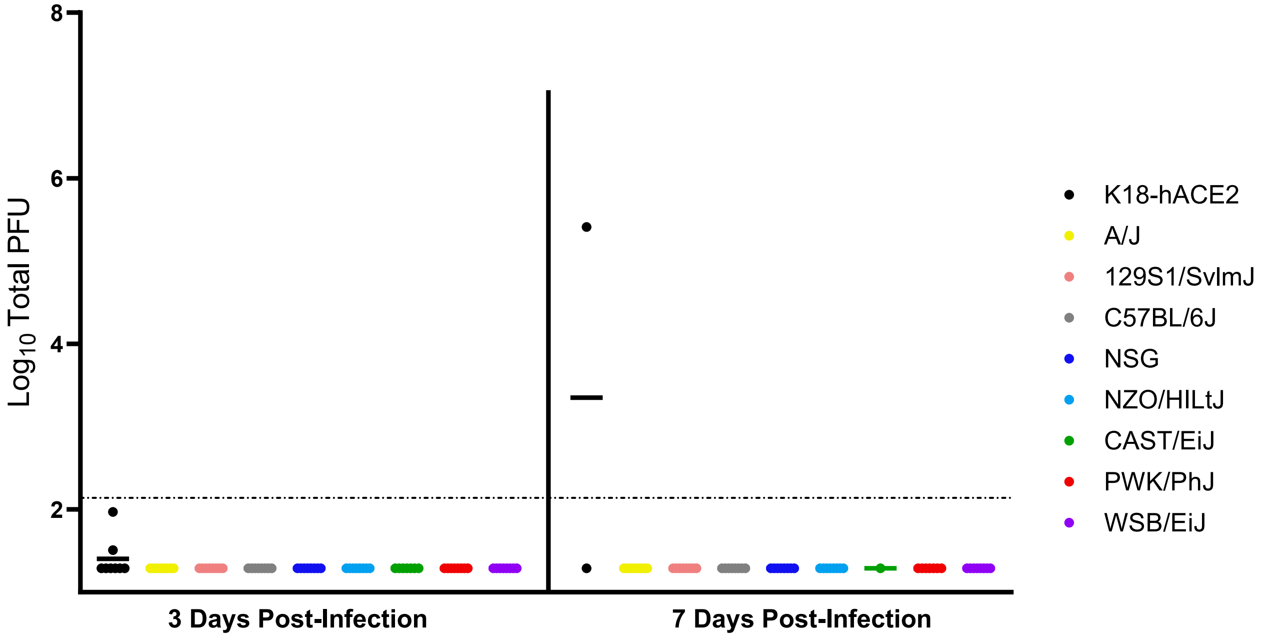


**Supplemental Figure 1. Analysis of infectious SARS-CoV-2 in brain tissue from the indicated mouse strains at days 3 and 7 post-infection.** Mice (n=8/strain per time point per virus, 4 males, 4 female) were infected with Beta SARS-CoV-2 variant and plaque assays were performed on brain homogenates (a).

a

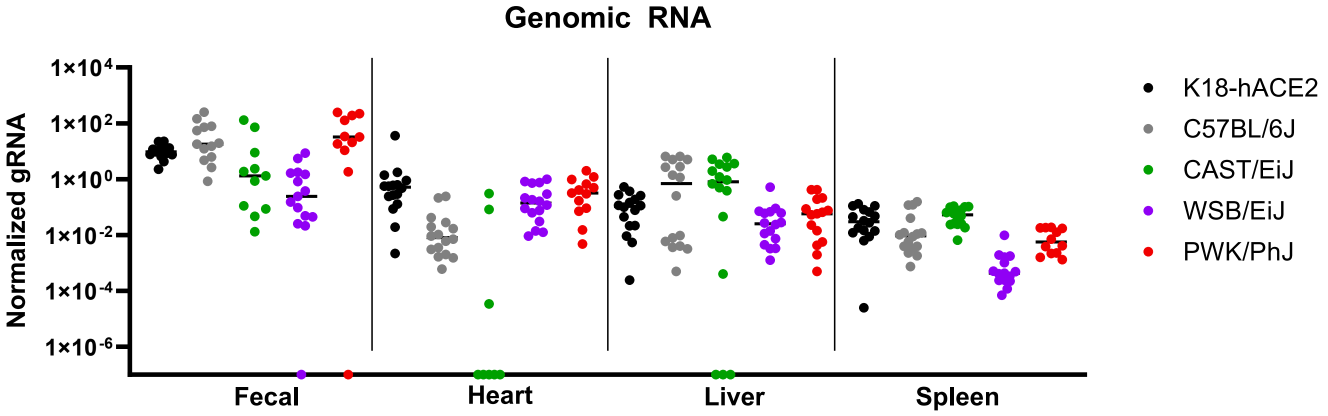


b

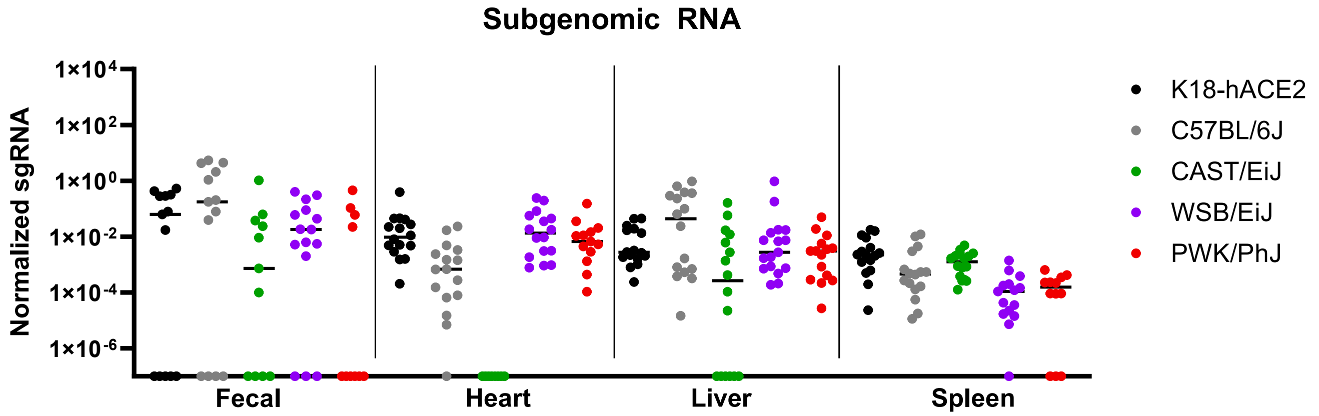


**Supplemental Figure 2. Analysis of viral genomic and subgenomic RNA levels in selected tissues and samples.** Each data point indicates a sample or tissue from an individual animal (n=16/strain per time point, 8 males, 8 female) for viral genomic RNA (a) or subgenomic RNA (b) levels. Viral RNA signals were normalized to endogenous mouse RNase P mRNA. Data points on the X axes were undetectable for viral RNA.

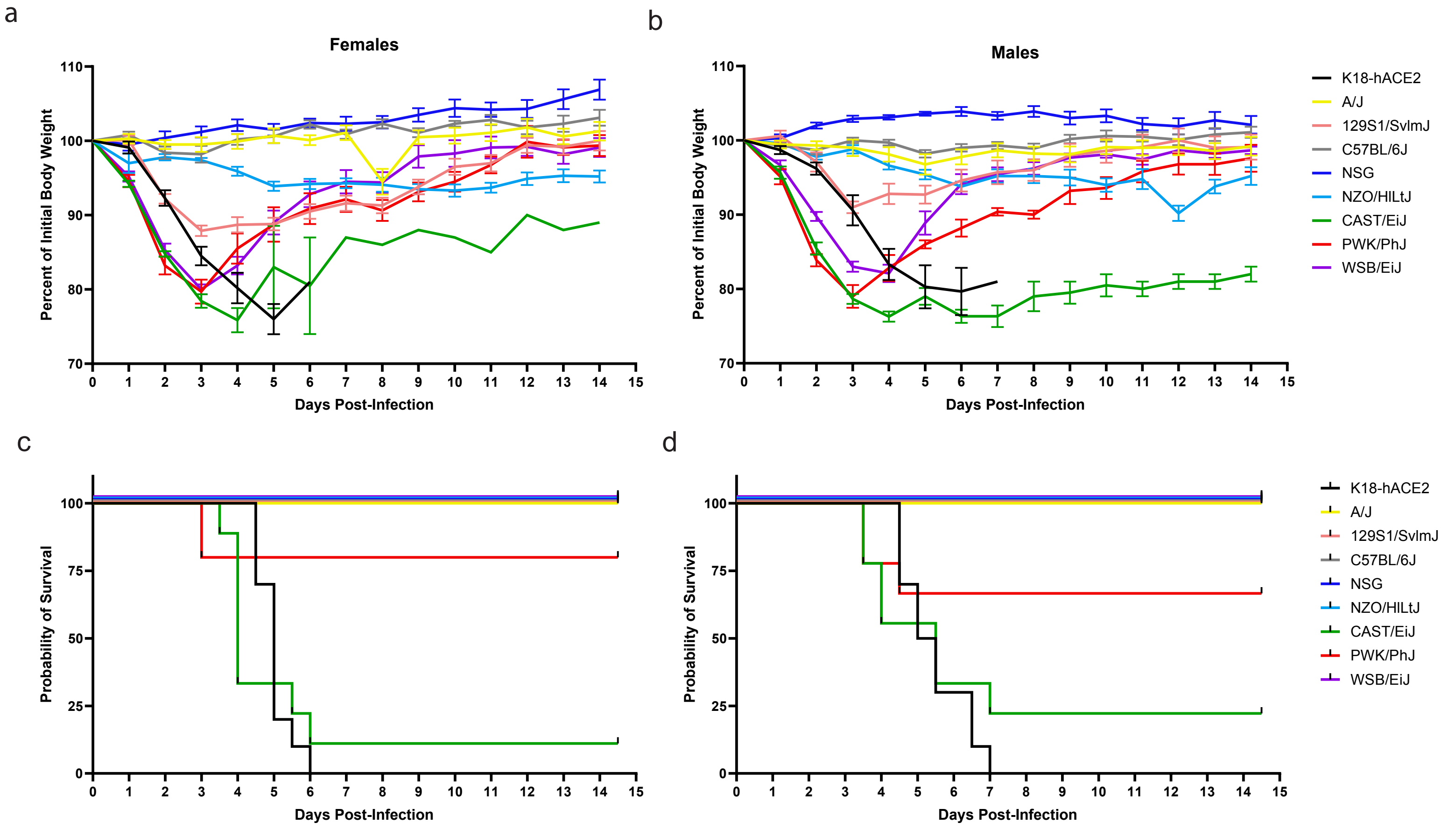


**Supplemental Figure 3. Evaluation of genetically diverse mouse strains for susceptibility to SARS-CoV-2 Gamma variant infection in comparison to transgenic K18-hACE2 mice.** Mice (n=20/strain, 10 males, 10 female) were infected with 1x105 PFU SARS-CoV-2 Gamma VOC. Female and male body weights (a-b) and were monitored daily for 14.5 days post-infection. Data are graphed as mean percent of initial body weight ± SEM. (c-d) Probability of survival is shown for each strain.

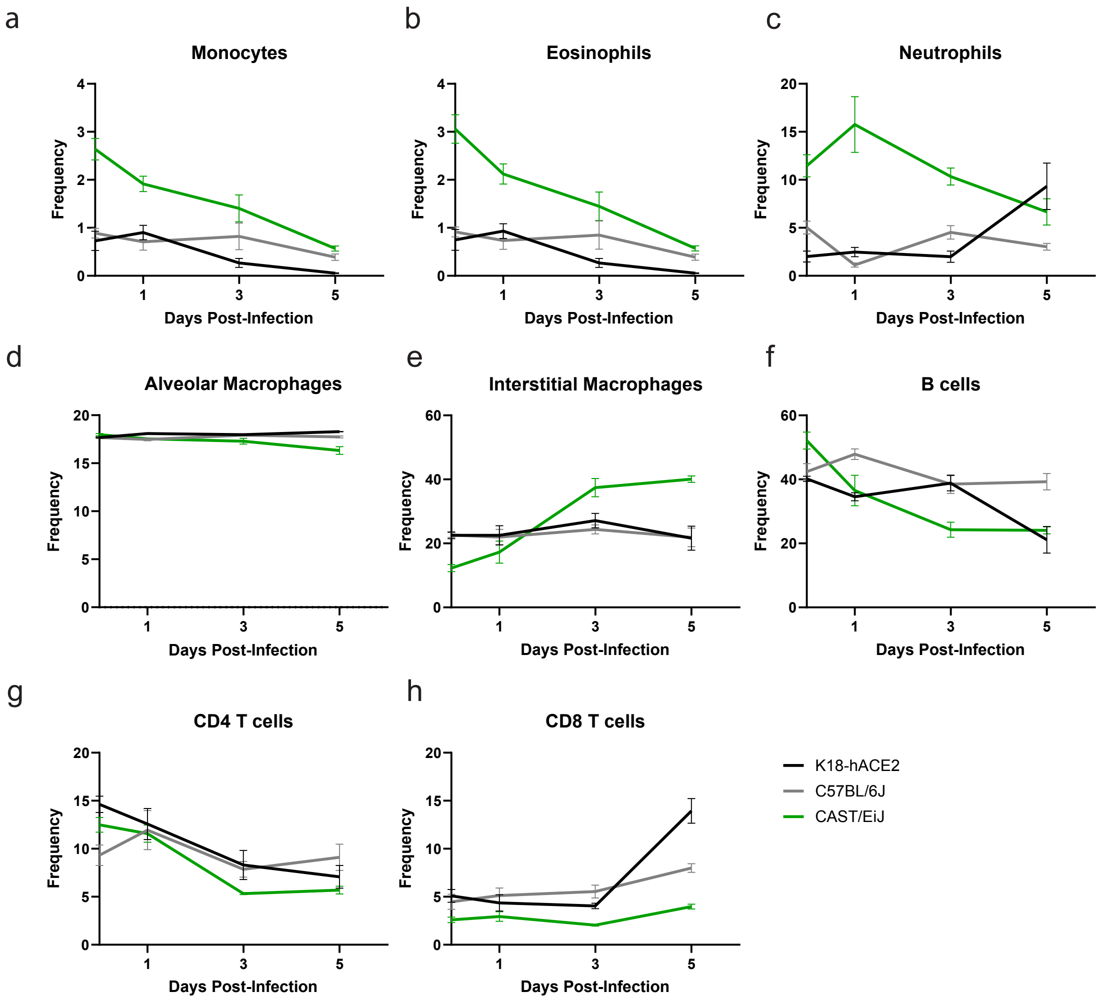


**Supplemental Figure 4. Flow cytometric analysis from SARS infected lung tissue show early indicators of survival.** Mice were infected with 1x105 PFU SARS-CoV-2 Beta VOC and lung tissue isolated for flow cytometric analysis. Frequencies in percent total of the indicated cell types is shown for each mouse strain at baseline (day 0) and at days 1, 3, and 5 post-infection (a-h). The data are representative of two experiments of similar design, wherein 5 mice were used per group.

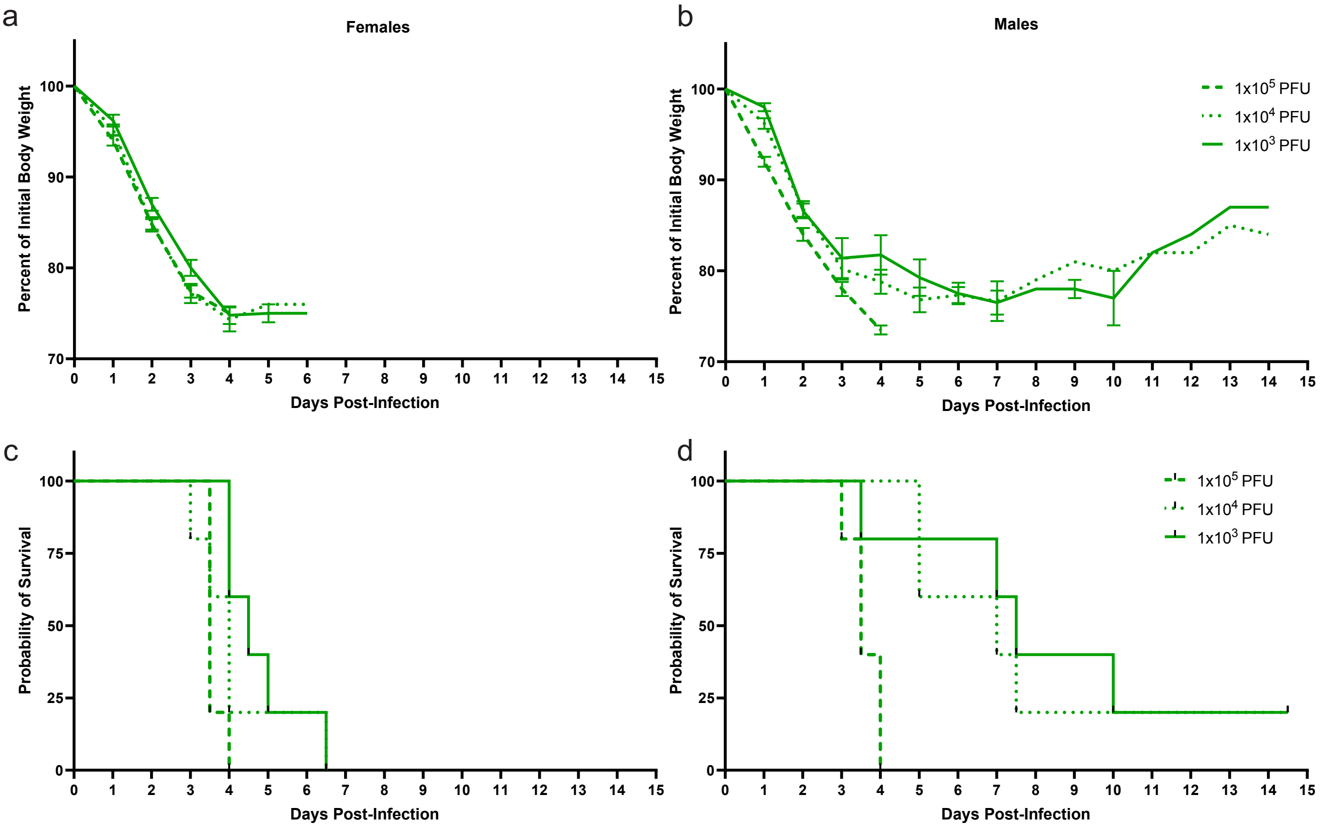


**Supplemental Figure 5. Analysis of CAST/EiJ female and male bodyweight and survival after challenge with SARS-CoV-2 Beta VOC doses.** Three different doses of SARS-CoV-2 Beta VOC were used to challenge female and male CAST/EiJ mice. Body weight (a) and survival (b) were monitored over 14.5 days. The data are representative of two experiments of similar design, wherein 10 mice (5 male and 5 female) were used per infectious dose.
